# Short Oxygen Pulses Enhance Creative Problem-Solving

**DOI:** 10.1101/2025.09.26.678411

**Authors:** Zenas C. Chao, Miyoko Street, Chien-Te Wu

**Author notes:** Correspondence: Zenas C. Chao.

## Abstract

Creativity is central to human innovation, yet it often fluctuates from moment to moment. Identifying simple interventions to reliably boost creativity has broad scientific and societal value. Here, we tested whether short-pulse oxygen inhalation enhances creative problem-solving. Sixty participants performed two established tasks: the Alternative Uses Test (AUT), capturing divergent idea generation, and the Fusion Innovation Test (FIT), assessing both divergent and convergent thinking. Oxygen (∼40% FiO_2_) was delivered in 1-minute pulses at 3–4-minute intervals, designed to align with intrinsic brain flexibility rhythms. Responses were scored for novelty, feasibility, and goal attainment using a validated GPT-based method. Linear mixed-effects regression revealed that oxygen significantly enhances both the quality and quantity of creative ideas across tasks. These findings demonstrate that a safe, low-cost physiological intervention can augment creative performance, providing a new link between oxygen metabolism, neural flexibility, and problem-solving.

**Significance statement:** Human progress depends on creativity, yet individuals often struggle to access their full creative potential. We demonstrate that brief cycles of enriched oxygen inhalation enhance creativity across distinct problem-solving tasks. This safe and low-cost protocol increased both originality and productivity, pointing to oxygen metabolism as a previously underappreciated driver of flexible thinking. The work introduces a novel, scalable approach to support creativity at a time when human innovation is essential alongside advances in artificial intelligence.

## Introduction

Creativity is increasingly vital in the era of artificial intelligence, where human originality and problem solving complement machine intelligence. Yet, effective strategies to enhance creativity remain limited. One promising avenue is the use of normobaric high-concentration oxygen, which has been shown to be safe under controlled conditions (FiO_2_ <60% for up to 24 hours) (1). Prior work demonstrates that creativity can be enhanced through cognitive training (2) and brain stimulation (3). Oxygen inhalation may offer a more accessible and economical alternative, either as a standalone intervention or in combination with these existing approaches.

To examine this possibility, we applied complementary assessments of divergent and convergent thinking. The Alternative Uses Test (AUT) requires participants to generate unconventional uses for everyday items (e.g., umbrella) (4). Responses are evaluated for novelty and feasibility using a validated GPT-based method that closely matches human judgments (5). While the AUT captures divergent idea generation, real-world creativity also requires convergent selection of effective solutions. To assess this process, we employed the Fusion Innovation Test (FIT), which asks participants to solve open-ended problems involving personal or societal goals by creatively combining unrelated elements. The FIT measures both divergent and convergent processes, with responses scored for novelty, feasibility, and goal attainment using automated GPT-based evaluation (6).

Our previous work decoding brain activity during AUT and FIT revealed intrinsic fluctuations in neural flexibility cycling at approximately 3.5 minutes (7). Building on this finding, we designed a novel oxygen delivery protocol using 1-minute short pulses in cycles of ∼3–4 minutes, aiming to entrain brain rhythms while reducing potential oxygen toxicity during long-term use. Using this protocol, we conducted a within-subject study with 60 participants and demonstrated that short-pulse oxygen inhalation significantly enhanced both the quality and quantity of creative performance across tasks.

## Results

We assessed creativity using two established tasks, the AUT and FIT. In each AUT trial, participants had 1.5 minutes to generate as many creative uses as possible for an everyday object (Figure 1A). In each FIT trial, participants had 4 minutes to combine two items to achieve a specified goal and produce as many creative solutions as possible (Figure 1B). AUT responses were scored for novelty and feasibility using a GPT-based automated evaluation method (5), and overall creativity was defined as their geometric mean. FIT responses were evaluated for novelty, feasibility, and goal attainment using a similar method (6), with overall creativity again defined as the geometric mean. These measures have been validated as reflecting real-world creativity (7). Fluency, defined as the total number of responses, was also recorded.

**Figure 1.**
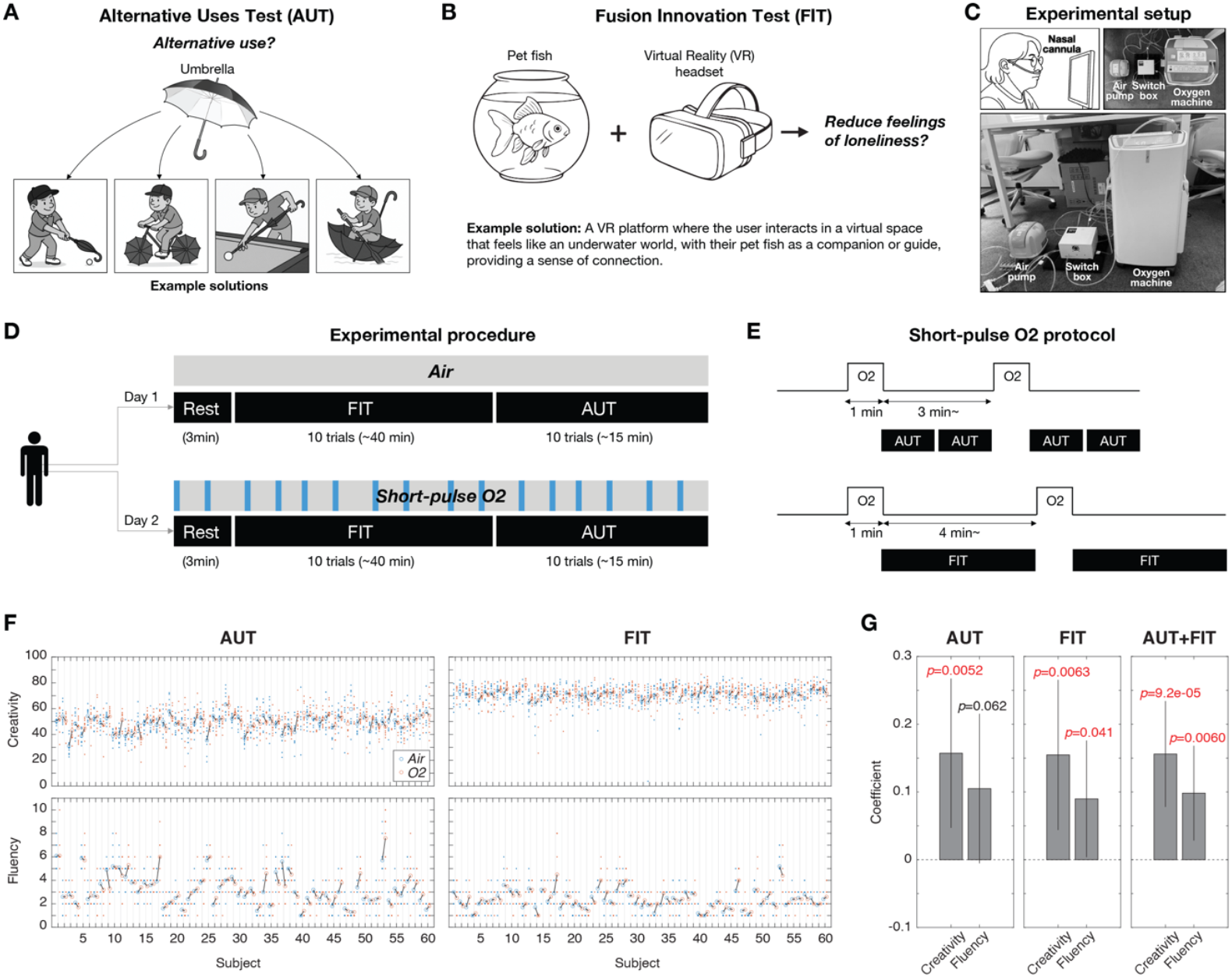
Oxygen enhancement effect on creativity performance. (**A**) Alternative Uses Test (AUT). The example item shown is “umbrella,” with possible creative solutions illustrated. In the actual test, both items and responses are presented in text only; figures here are for illustration. (**B**) Fusion Innovation Test (FIT). Two sample question items and a goal are presented, along with an example solution. As with AUT, the real test uses text only. (**C**) Experimental setup for oxygen delivery. (**D**) Experiential procedure for within-subject design. (**E**) Oxygen protocol for AUT and FIT sessions. (**F**) Creativity scores and fluency for each subject in AUT and FIT. Each dot represents the average creativity across multiple solutions and the fluency of a single trial (blue: *Air*, red: *O2*). Circles indicate the mean across 10 trials, with black lines denoting the difference between conditions. (**G**) Regression beta coefficients with 95% confidence intervals for AUT, FIT, and combined data (AUT+FIT). p-values are indicated, with significant values highlighted in red. Images in panels A and B were generated with ChatGPT (OpenAI).

Data were collected from 60 participants (43 males, 17 females; age range 19–31 years; mean ± SD: 22.7 ± 2.3 years). Each participant completed 20 AUT and 20 FIT questions across two experimental days. On one day, participants received normal air (*Air* condition), and on the other, they received short-pulse oxygen at ∼40% inspired fraction (*O2* condition) delivered via nasal cannula (Figure 1C). The assignment of condition to day was randomized across participants. Each day, participants completed one block of 10 AUT questions and one block of 10 FIT questions. Task order (AUT or FIT) was counterbalanced across participants but fixed within individuals (Figure 1D). Short-pulse oxygen was delivered for 1 minute, followed by a 3-minute interval during the AUT (covering two questions) and a 4-minute interval during the FIT (covering one question) (Figure 1E). During the intervals, normal air was supplied at the same flow rate.

For each question, creativity scores were averaged across solutions, and the mean creativity and fluency across all questions and participants are shown in Figure 1F. To assess oxygen effects within individuals, mean creativity and fluency scores from 20 AUT and 20 FIT questions were z-scored within each task for each participant. These normalized scores were analyzed using a linear mixed-effects regression, with condition (*Air* vs. *O2*) as a fixed effect and random intercepts for participant, question, and day nested within participant.

In the AUT, condition (*O2* vs. *Air*) significantly increased creativity (β = 0.16, SE = 0.06, 95% CI [0.05, 0.27], t(1192) = 2.80, p = 0.005) and showed a marginal increase in fluency (β = 0.11, SE = 0.06, 95% CI [–0.01, 0.22], t(1198) = 1.87, p = 0.062) (Figure 1G, left). In the FIT, oxygen significantly enhanced both creativity (β = 0.15, SE = 0.06, 95% CI [0.04, 0.27], t(1179) = 2.74, p = 0.006) and fluency (β = 0.09, SE = 0.04, 95% CI [0.00, 0.18], t(1198) = 2.05, p = 0.041) (Figure 1G, middle). When combining AUT and FIT, oxygen significantly improved both creativity (β = 0.16, SE = 0.04, 95% CI [0.08, 0.23], t(2372) = 3.92, p = 9.2e-5) and fluency (β = 0.10, SE = 0.04, 95% CI [0.03, 0.17], t(2397) = 2.75, p = 0.006) (Figure 1G, right). Full statistical results are provided in Table 1. Together, these results indicate that short-pulse oxygen enhanced both the quality and quantity of generated solutions.

**Table 1:** Summary of linear mixed-effects models.

| Task | Outcome | $\beta$<br>(O<sub>2</sub> vs Air) | SE | 95% CI | t(df) | p | N<br>(trials) | Residual<br>SD | Random:<br>Question<br>SD | AIC | BIC | logLik |
| --- | --- | --- | --- | --- | --- | --- | --- | --- | --- | --- | --- | --- |
| AUT | Creativity | 0.16 | 0.06 | [0.05,<br>0.27] | 2.80(1192) | 0.005 | 1194 | 0.53 | 0.81 | 3331.1 | 3361.6 | −1659.5 |
|  | Fluency | 0.11 | 0.06 | [−0.01,<br>0.22] | 1.87(1198) | 0.062 | 1200 | 0.53 | 0.82 | 3357.1 | 3387.6 | −1672.6 |
| FIT | Creativity | 0.15 | 0.06 | [0.04,<br>0.27] | 2.74(1179) | 0.006 | 1181 | 0.97 | ~0 (negl.) | 3294.5 | 3324.9 | −1641.2 |
|  | Fluency | 0.09 | 0.04 | [0.00,<br>0.18] | 2.05(1198) | 0.041 | 1200 | 0.73 | 0.22 | 2761.3 | 2791.8 | −1374.7 |
| AUT+FIT | Creativity | 0.16 | 0.04 | [0.08,<br>0.23] | 3.92(2372) | 9.2e−5 | 2375 | 0.94 | 0.25 | 6613.5 | 6653.9 | −3299.7 |
|  | Fluency | 0.10 | 0.04 | [0.03,<br>0.17] | 2.75(2397) | 0.006 | 2400 | 0.86 | 0.14 | 6181.4 | 6221.9 | −3083.7 |

## Discussion

Our findings demonstrate that short pulses of enriched oxygen reliably enhance a general capacity for flexible thinking, rather than promoting task-specific strategies. We propose two complementary mechanisms that may underlie this effect. One possibility is that intermittent oxygen delivery aligns with intrinsic neural rhythms of flexibility (7), entraining and amplifying these oscillations to broaden the brain’s accessible state space. Another possibility is what we call the oxygen massage hypothesis. Our prior fMRI work revealed that the impact of oxygen on resting-state activity and connectivity varies across regions, allowing us to construct a brain oxygen sensitivity topography (BOST) (8). This map showed that the onset and offset of oxygen inhalation produce prolonged activation within the default mode network (DMN), a system thought to support creativity and cognitive flexibility (9, 10). We hypothesize that the brain must rapidly adapt to fluctuations in blood oxygen (both increases and decreases) such that short oxygen pulses “force” adaptive responses. In this sense, pulsed oxygen may act like passive exercise or a massage, promoting neural flexibility. The choice of a 1-min pulse was also informed by the BOST study, where brain-wide BOLD responses increased and plateaued after approximately 45 seconds of high-concentration oxygen.

Physiological evidence provides further support for why oxygen may exert these effects. Under hyperoxic conditions, lactate levels decrease while glucose remains stable (11), suggesting a shift from aerobic glycolysis toward oxidative metabolism that produces excess energy. The DMN appears particularly sensitive to such metabolic changes, as both hypoxia and altered glucose availability have been shown to modulate its activity and connectivity (12, 13). This metabolic profile may explain why oxygen pulses preferentially influence the DMN and, in turn, cognitive flexibility.

Several limitations should be acknowledged. Parameters of oxygen delivery (such as duration, interval, concentration, and pressure) remain to be optimized for maximal benefit. Additional behavioral assessments, including real-world problem-solving tasks in naturalistic environments, will be necessary to validate the ecological impact of this intervention. Our study focused on healthy young adults in controlled laboratory settings, leaving open the question of whether similar effects would be observed in other populations, such as children, older adults, or clinical groups. Finally, although our pulsed protocol was designed to minimize toxicity risk, further work is needed to establish optimal dosing schedules and assess potential cumulative effects.

## Methods

### Participants

Data were collected from 60 participants (43 males, 17 females), aged 19∼31 years (mean ± standard deviation: 22.7 ± 2.3 years) between July 2024 and April 2025. Most were university students, and all were native Japanese speakers, with no history of neurological disorders and respiratory disorders. Prior to the experiment, participants provided written informed consent. The study was approved by the Research Ethics Committee of the University of Tokyo (No. 24-027). Participants were recruited via a website (https://www.jikken-baito.com).

### Oxygen delivery

Air and oxygen were delivered by an air pump (GEX AIR PUMP e-AIR 9000FB) and a medical oxygen concentrator (LiteTEC-5B, JMDN12873002; Figure 1C), respectively, at a flow rate of 5 L/min through a nasal tube (OX-20, Atom Medical Corp). The concentrator supplied ∼90% oxygen under normobaric conditions. Switching between air and oxygen was automated in MATLAB and controlled via a custom switch box developed by Daikin Industries, Ltd. (Figure 1C).

### Experimental setup and procedure

Data were collected in a quiet, well-lit room maintained at 25 °C and 50% relative humidity. Participants sat in front of a 43-inch monitor for the presentation of questions and instructions and entered their responses into a Google Form on a tablet with a keyboard.

Each participant completed 20 AUT questions and 20 FIT questions across two experimental days. Each day consisted of one block of 10 AUT questions and one block of 10 FIT questions (5 personal-improvement goals and 5 social-developmental goals, the two FIT categories). Block order was reversed on the second day, and the starting task (AUT or FIT) was counterbalanced across participants.

At the beginning of each block, a 3-minute resting period was included, during which participants fixated on a central cross while breathing normal air. Short-pulse oxygen or air was then delivered for 1 minute, followed by a 3-minute interval to answer two AUT questions or a 4-minute interval to answer one FIT question, during which only normal air was delivered. On the first day, participants completed one practice trial to familiarize themselves with the procedure.

For each trial, participants read an AUT or FIT question on the monitor and entered as many responses as possible within the time limit (1.5 min for AUT, 4 min for FIT). They then advanced to the next question. Responses left unfinished when the time expired were excluded from analysis.

### Response evaluation and regularization

Participants’ responses were evaluated automatically using GPT-4o, with prompts adapted from previous studies (AUT: Kern et al., 2024; FIT: Wu et al., 2024). For AUT, each response was rated on two dimensions: Novelty and Feasibility. For FIT, each response was rated on three dimensions: Novelty of item combinations, Feasibility of item combinations, and Goal Attainment. All scores were scaled from 1 to 100. For AUT, overall creativity was defined as the geometric mean of *N* and *F* (√(*N* × *F*)). For FIT, overall creativity was defined as the geometric mean of *N, F*, and *G* (^3^√(*N* × *F* × *G*)).

### Regression analysis

We analyzed the effects of oxygen on *Creativity* and *Fluency* using mixed-effects regression models implemented in MATLAB (glmfit.m). For each task separately (AUT and FIT), creativity scores were modeled as a function of *Condition* (*O2* vs. *Air*) with random intercepts for *Participant, Participant:Question*, and *Participant:Day*:

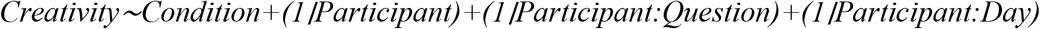

Fluency was analyzed with an identical structure:

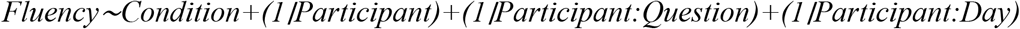

To test effects across both tasks jointly, *Condition* and *Task* (AUT vs. FIT) were included as fixed effects, with the same random effects structure:

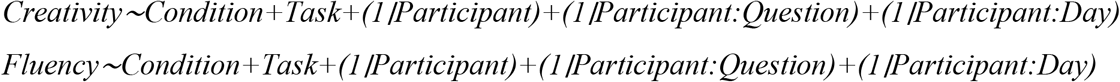

Including random intercepts for *Participant* accounted for individual differences in overall performance, while the *Participant:Question* and *Participant:Day* terms captured variability associated with specific question items and day-to-day fluctuations.

## Author contributions

Z.C.C. conceptualized the study. C.T.W. implemented the experiment. M.S. collected the data. Z.C.C. performed data analysis and wrote the paper. M.S. edited the paper. All authors contributed to and approved the final paper.

## Acknowledgments

We thank Dr. Iver Hsieh for developing and maintaining the web-based task interface, Dr. Yu-Shian Su for acquiring the GPT scores, and Dr. Yan Zhang for experimental support. This work was supported by the World Premier International Research Center Initiative (WPI), MEXT, Japan (to Z.C.C.), and the IRCN–Daikin SCP (to Z.C.C.).

